# Dynamic expression of tRNA-derived small RNAs define cellular states

**DOI:** 10.1101/158501

**Authors:** Daniel GR Yim, Srikar Krishna, Vairavan Lakshmanan, Judice LY Koh, Jung Eun Park, Jit Kong Cheong, Joo Leng Low, Michelle JS Lim, IP Junyu, Jie Min Nah, Iain BH Tan, N Gopalakrishna Iyer, Huili Guo, Siu Kwan Sze, Srikala Raghavan, Dasaradhi Palakodeti, Ramanuj Dasgupta

**Author notes:** Co-first author (Daniel GR Yim, Srikar Krishna, Vairavan Lakshmanan).

## Abstract

Transfer RNA (tRNA)-derived small RNAs (tsRNAs) have recently emerged as important regulators of protein translation and shown to have diverse biological functions. However, the underlying cellular and molecular mechanisms of tsRNA function in the context of dynamic cell-state transitions remain unclear. Here we report the identification of a set of tsRNAs upregulated in differentiating mouse embryonic stem cells (mESCs). Mechanistic analyses revealed primary functions of tsRNAs in regulating polysome assembly and translation. Notably, interactome studies with differentially-enriched tsRNAs revealed a switch in associations with ‘effector’ RNPs and ‘target’ mRNAs in different cell-states. We also demonstrate that a specific pool of tsRNAs can interact with Igf2bp1, an RNA-binding protein, to influence the expression of the pluripotency-promoting factor-c-Myc, thereby providing evidence for tsRNAs in modulating stem cell-states in mESCs. Finally, tsRNA expression analyses in distinct, heterologous cell and tissue models of stem/transformed versus differentiated/normal states reveal that tsRNA-mediated regulation of protein translation may represent a global biological phenomenon associated with cell-state transitions.

**One Sentence Summary:** Identification and functional characterization of tRNA-derived small RNAs (tsRNAs) in cell state switches.

## Introduction

Cell differentiation is a highly dynamic process involving the spatial-temporal expression of proteins that control specific cellular phenotypes. Elucidating the molecular mechanisms underlying cell-state transitions is important in understanding the process of differentiation. Gene expression is tightly regulated at the transcriptional, post-transcriptional and translational levels. Multiple studies have identified the function of miRNAs in modulating gene expression driving pluripotency and differentiation (Ivey and Srivastava, 2010). However, miRNAs represent only a subset of non-coding RNAs (ncRNAs); other small ncRNAs as modulators of pluripotency/differentiation remains relatively under-explored.

Transfer RNA (tRNA)-derived small RNAs (tsRNAs) are a class of ncRNAs first identified as regulators of *Tetrahymena* stress response (Lee SR, et al. 2005). Recently, species of tsRNAs processed from different parts of the parent/precursor tRNA have been identified (Anderson P and Ivanov P, 2014; Keam SP, et al. 2015) and shown to have roles in biological processes such as tumor suppression (Goodarzi et al., 2015) and paternal epigenetic inheritance (Chen Q et al., 2016; Sharma et al., 2016). Interestingly, a specific class of tsRNAs derived from the 5’-half of tRNAs have been implicated in translational inhibition (Ivanov et al., 2011; Yamasaki et al., 2009, Sobala A, et al. 2013).

Here, we report an enrichment of tsRNAs derived from the 5’-half of tRNAs during retinoic acid (RA)-induced differentiation of mouse embryonic stem cells (mESCs). Functional characterization using overexpression and knockdown studies suggest that tsRNAs can modulate mESC differentiation. Polysome profiles during RA-induced differentiation inversely correlated with tsRNA expression, suggesting a role for tsRNAs in modulating translation during differentiation. Small RNA sequencing from isolated polysome fractions revealed significantly increased association of tsRNAs with monosome (80S) and polysome units. These results suggest an additional role for tsRNAs in modulating active translation of mRNAs in addition to the previously reported regulation at translational initiation (Ivanov P, et al. 2011). Intriguingly, the RNA and proteome interaction studies with tsRNA tsGlnCTG revealed a hitherto unknown role in regulating cell states and lineage specification by modulating translation of specific transcripts by virtue of their interaction with different subsets of RNA binding proteins. Specifically, we provide a mechanism for tsRNAs-mediated regulation of c-Myc through its interaction with Igf2bp1, an RNA-binding protein that stabilizes c-Myc mRNA (Weidensdorfer et al., 2009). Additionally, we also observed similar enrichment of tsRNAs in a broad range of heterologous differentiation cell states, including patient-derived cancer cells, underscoring the potential role for tsRNAs as generic modulators of cell differentiation. Taken together our data suggests a fundamental function for tsRNAs in regulating translation of mRNAs that are critical to cell-state transitions.

## Results

### Differentially expressed tsRNAs modulate stem cell pluripotency

Deep sequencing of small RNAs in mESCs cultured under different conditions including Wnt3a+LIF (enhanced pluripotency), LIF alone (baseline pluripotency), and retinoic acid (RA) (differentiation) (Fig. S1A) revealed a distinct population of 30-35 nt species enriched during differentiation (done in biological duplicates) (Fig. 1A). Intriguingly a large proportion of these 30-35 nt species mapped to tRNAs, and exhibited dynamic expression between stem and differentiating states. Specifically, in RA-treated differentiating mESCs 81.4% of these reads mapped to tRNAs, which dropped to 72% in LIF and to 50.3% in Wnt3a+LIF culture conditions (Fig. 1B and Table S1). These results suggest that the expression of tsRNAs inversely correlate with the pluripotent state of mESCs. Notably, most tsRNAs were derived from the 5’ halves, starting from nucleotide 1-4 of parent tRNAs and terminating prior to the anticodon loop (Figs. 1C, 1D and S1B), suggesting specific processing of tRNAs. The tsRNAs enriched during RA-induced differentiation predominantly correspond to GlnCTG, GlyGCC, GluTTC, and ValAAC tRNAs (Table S2), and were validated by small RNA qPCR (Fig. 1E). Since we observed tsRNA enrichment during differentiation, we hypothesized that the change in the tsRNA levels might be crucial for stem cell state transitions. To this front, we performed alkaline phosphatase (AP) assays as a read-out for pluripotency while overexpressing (using tsRNA mimics) or knocking down (using antisense oligonucleotides, ASOs) tsRNAs. Overexpression of these tsRNAs, individually or collectively, inhibited AP activity in pluripotent mESCs (Fig. 1F), with a concomitant increase in the expression of specific differentiation markers (Fig. S1C). Conversely, inhibition of tsRNAs with pooled antisense oligonucleotides (ASOs) increased the AP activity in RA-treated mESCs (Fig. 1G), which was further confirmed by the increase in expression of pluripotency markers (Fig. S1D). Taken together, these data reveal that the tsRNA-dependent modulation of cell fate appears to be subtle but significant, suggesting that tsRNAs may function as fine-tuners of biological processes, akin to microRNAs (miRNAs), reviewed in (Lai, 2015).

**Figure. 1.**
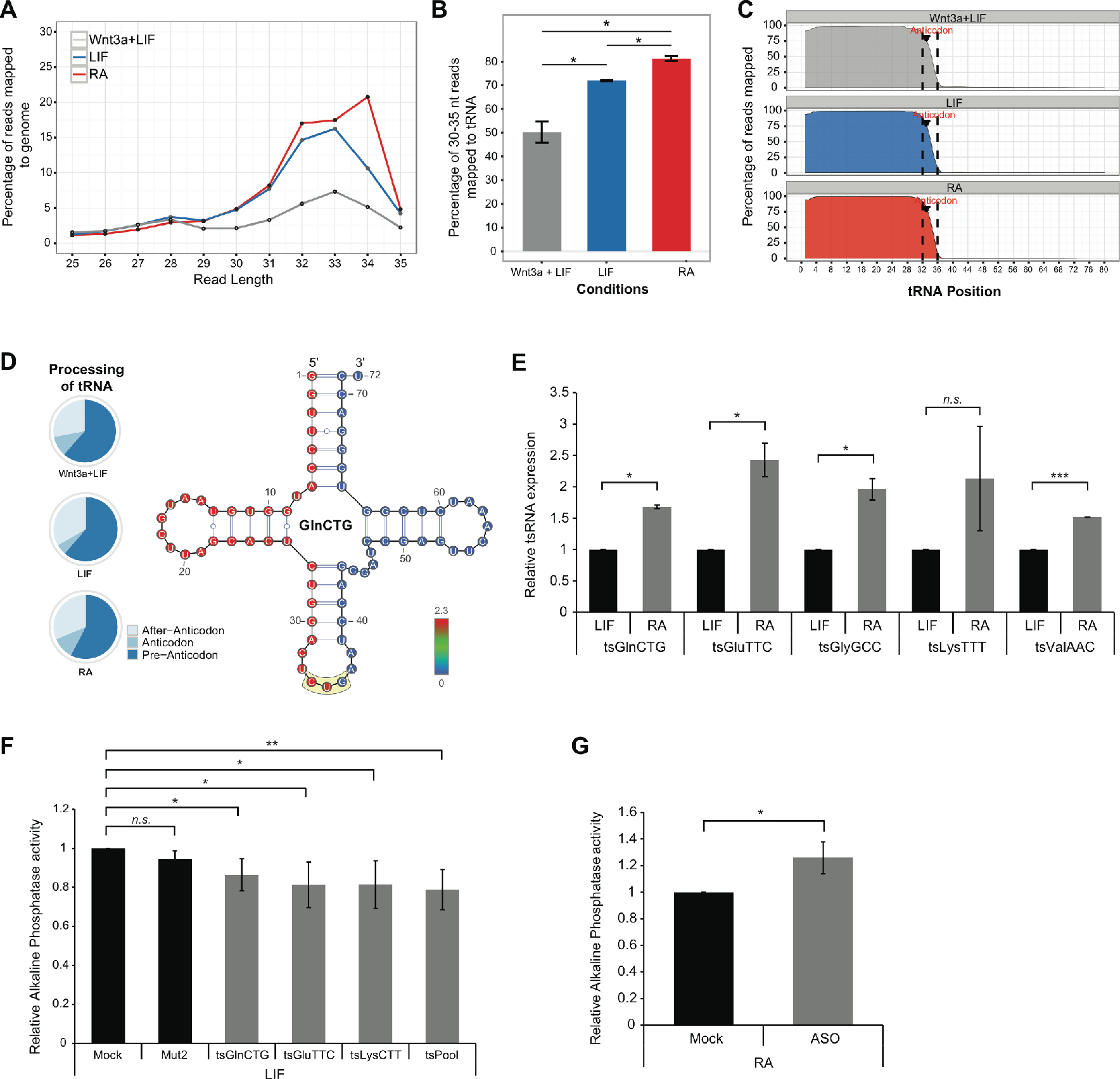
Expression and functional characterization of tsRNAs in mESC. **(A)** Distribution of 25-35 mer reads mapping to the mouse genome. **(B)** Proportion of 30-35 nt genomic reads matching mouse tRNA sequences. (Error bars represent SEM, significance was calculated using unpaired t-test) **(C)** Perbase sequence coverage of 30-35 nt reads across parental tRNA sequence. Anticodon positions are demarcated by discontinuous lines (Blue = LIF-treated, Grey = Wnt3+LIF-treated, Red = RA-treated mESCs). **(D)** *Left:* Distribution of read ends to regions indicated on parental tRNAs. *Right:* Positional heat map of reads mapped to tRNA^GlnCTG^ residues. **(E)** Expression of top RA-enriched tsRNAs quantified with small RNA qPCR (Error bars represent SD, significance was calculated using unpaired t-test). **(F)** Effect of overexpression of tsRNA mimics on mESC pluripotency (grown in LIF condition) as measured by alkaline phosphatase activity (Error bars represent SD, significance was calculated using unpaired t-test). **(G)** Effect of ASO-mediated inhibition of tsRNAs on alkaline phosphatase activity in RA-treated mESCs (Error bars represent SD, Significance was calculated using unpaired t-test). *, *P* value < 0.05, **, *P* value < 0.01, *n.s.*, not significant.

### Dynamic tsRNA expression regulates protein translation and polysome assembly

Translational regulation plays a pivotal role in the temporal regulation of genes critical for transition of cells from pluripotency to differentiated states (Sampath et al., 2008). Since tsRNAs are known to regulate protein translation (Ivanov P, et al., 2011; Yamasaki et al., 2009), we hypothesized that tsRNA abundance would be anti-phasic with protein synthesis activity. To test this hypothesis, we measured the levels of tsRNAs and translation at different time points during differentiation.

Given that most of the 30-35 nt RNAs in mESCs were tsRNAs, we used 30-35 nt RNAs measured using Bioanalyzer electrophoresis as a proxy for the levels of tsRNAs upon differentiation. In comparison to LIF, RA-treated mESCs showed reduced levels of tsRNAs after 12 hours, reaching a maximum at ∼48 hours, and then gradually diminishing to a second minimum 6 days post-RA treatment (Fig. S2A). In addition, we measured the polysome to monosome ratio (P:M), an indicator of active translation (Chassé et al., 2017) (Fig. 2A) in LIF- and RA-treated mESCs. Local non-linear regression of tsRNA expression and P:M ratios at each time-point of differentiation revealed a potential anti-phasic relationship between active translation with the levels of tsRNAs (Fig. 2B). Protein translation in mESCs assayed by L-azidohomoalanine (AHA) incorporation further validated the same overall trend of translation (Fig. 2C and 2C’). The anti-phasic nature of tsRNA expression and protein translation suggests a role of tsRNA in translational regulation during RA-mediated differentiation. To test this, we utilized the *in vitro* biochemically-defined, rabbit reticulocyte lysate system to measure general translation activity in the presence of exogenously added tsRNAs. Indeed, we observed dose-dependent inhibition of protein translation upon the addition of a tsRNA-mimic for tsGlnCTG (Fig. 2D), suggesting that endogenous changes in total tsRNA levels may regulate translation of nascent proteins. To gain mechanistic insights into the effect of tsRNAs on translation, we used electron microscopy to observe the impact of tsRNA overexpression on polysome structures in rabbit-reticulocyte lysates. Upon the addition of mRNA to the lysates, the ribosomes aligned into string-of-beads-like structures (polysomes) (Fig. S2C, left). This assembly was markedly abrogated with the addition of tsGlnCTG mimic (Fig. S2C, middle), which formed undefined clusters resembling stress granules (SG). This observation is in agreement with previous reports that tsRNAs localizes to SGs in response to cellular stress signals (Emara et al., 2010; Yamasaki et al., 2009). It is important to note that the effect of tsRNAs in forming these SG-like clusters was dependent on the presence of mRNA (Fig. S2C, right). Therefore, tsRNAs may serve as molecular brakes for polysome assembly with consequent translational repression.

**Figure. 2.**
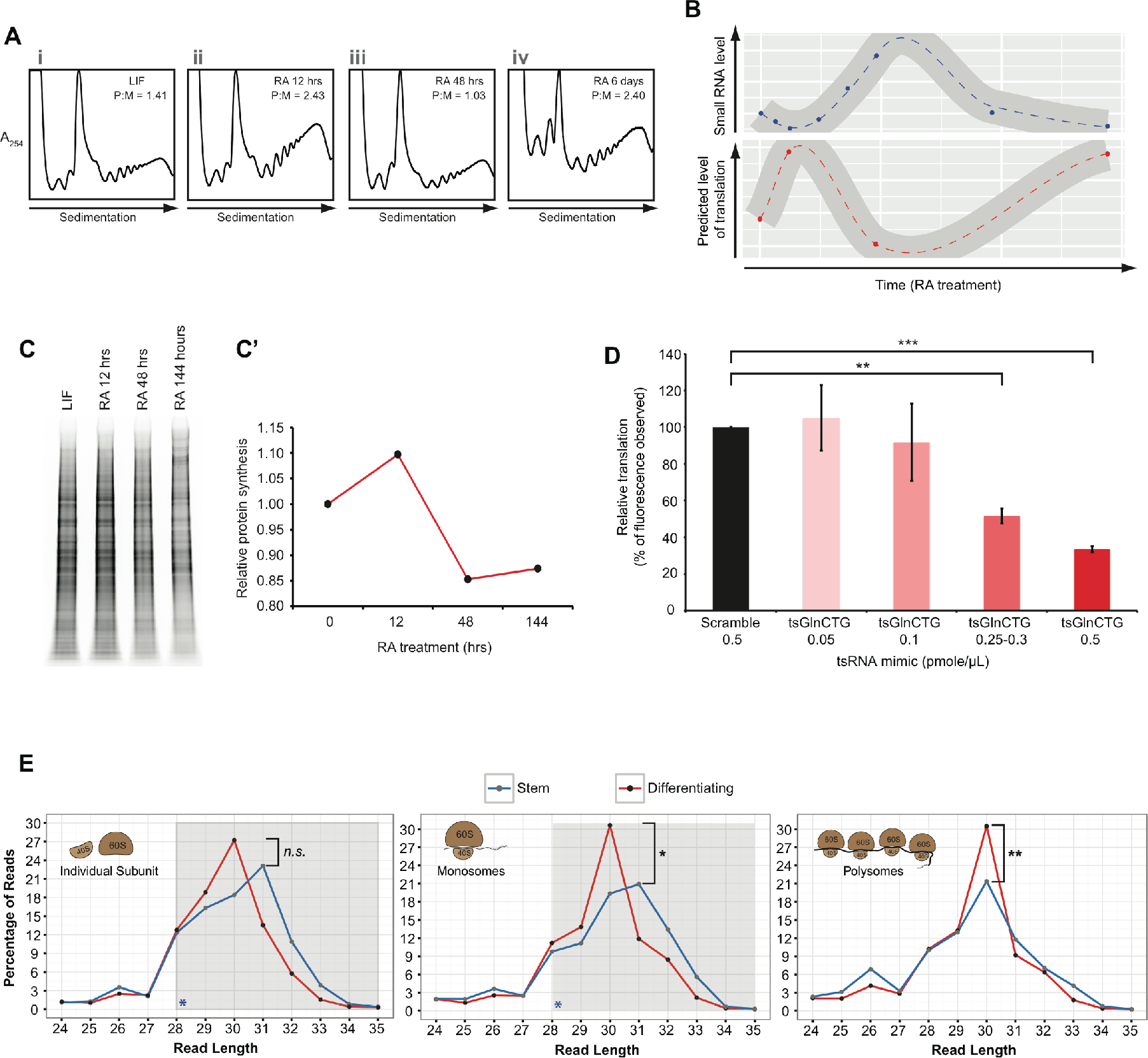
Molecular function of tsRNAs. **(A)** Polysome profiles and corresponding Polysome:Monosome (P:M) ratios at representative RA time-points shown in Fig. S3A (Points i to iv). **(B)** Loess curve-fitting modeling of changes in small RNA levels and predicted translation activity over a time-course of RA differentiation. Darker grey areas indicate standard error distribution of the regression. **(C and C’)** *In vivo* translation of nascent proteins. SDS-PAGE image of AHA-incorporated proteins at indicated cell states (C), and relative TAMRA signal (C’) normalized against corresponding levels of total protein (Fig. S2A). **(D)** Effects of tsGlnCTG on *in vitro* translation (Error bars represent SD, significance was calculated using unpaired t-test). **(E)** tsRNA distribution among the polysome fractions from indicated cell states. Shaded region highlights the change in the distribution of tsRNA pool (Calculated using change point analysis). *, *P* value < 0.05, **, *P* value < 0.01, ***, *P* value < 0.001.

Based on our observations that tsRNAs disrupt polysome structures and inhibit translation, next we investigated the association of tsRNAs with individual ribosomal subunits (40S, 60S), monosomes (80S) and polysomes in mESCs grown under stem or differentiating conditions (Fig. S2D). The tsRNA profiling (Table S1) revealed two interesting observations. First, we observed a significant shift in tsRNA profiles (change point analysis) sampled from individual subunits (40S and 60S) and monosomes (80S) during the differentiation (Fig. 2E, left and middle), and second, upon comparison of the peak maxima from the tsRNA profiles, we observed an enrichment of a 30-mer population (Wilcox’s test, *P* value < 0.05) in monosomes (80S) and polysomes derived from differentiating mESCs (Fig. 2E right). Taken together our data suggests a previously unreported function for tsRNAs in metazoan translation by associating with polysomes, in addition to their purported role in translation initiation via association with monosomes (Gebetsberger et al., 2012; Ivanov P, et al., 2011).

### tsRNAs regulate specific targets through their protein interactome

The enrichment of specific tsRNAs observed upon differentiation suggests a mechanism for specificity in target selection critical for this process. We explored this specificity by performing pull-down assays using a Biotin-tagged tsGlnCTG mimic, in mESCs treated with LIF or RA for 48hrs, which identified mRNA targets (Fig. S3A). Transcriptome analysis of tsGlnCTG pull-down revealed distinct association of transcripts (P value < 0.05) to the tsRNA in stem-(LIF) versus differentiating (RA) cell states (Fig. 3A). We observed 582 transcripts that associated equally with tsGlnCTG in both stem and differentiating states (Table S3). GO annotations of these transcripts revealed enrichments of genes involved in many basic cellular processes such as metabolism, and stress response (Table S4). In addition, tsGlnCTG showed enriched associations with 420 transcripts in RA condition versus 27 transcripts in the stem state, underscoring a potential functional role for tsGlnCTG in differentiation (Table S3). To understand the regulatory effect of tsRNAs on the target mRNAs, we investigated the binding preference of tsGlnCTG towards either ‘differentiation-responsive’ or ‘pluripotency-associated’ genes in both stem and differentiating conditions (see methods, section “tsRNA mediated mRNA pulldown library and analysis”). Of the 27 tsGlnCTG-associated transcripts that showed enrichment in the stem state, a preference was observed towards ‘differentiation-responsive transcripts’ (5 transcripts) such as *Cdh2*, compared to ‘pluripotency-associated transcripts’ (2 transcripts) (Fig. 3B and Table S3). Interestingly, upon RA mediated mESC differentiation, there was a marked increase (from 2 to 37) in the number of ‘pluripotency-associated’ transcripts associated with tsGlnCTG compared to the stem state. Some of these transcripts include *Klf4, Ina, Nr0b1* and *Lrrc2*, which are known pluripotency markers critical for the maintenance of the stem state (Sharova et al., 2007). However, we also observed an increase in tsGlnCTG association with pro-differentiation transcripts like *Myl3, CD44* and *Dkk3* (Cohen-Haguenauer et al., 1989; Quintanilla et al., 2014; Wang et al., 2015) (Fig. 3B’ and Table S3). While initially counter intuitive, upon closer examination of those transcripts, we noticed significant association towards genes involved in skeletal, cardiovascular, and blood vessel development compared to neuronal lineage (Table S4). Together, these data suggest that during neural lineage differentiation triggered by RA (Simandi et al., 2015), tsGlnCTG may gain an additional function in targeting and regulating the stability of transcripts driving non-neuronal pathways thus facilitating neuronal differentiation apart from regulating basic cellular processes. Furthermore, retrospective analysis of Brachyury expression (a neural mesodermal marker) revealed that it was unchanged upon ASO-mediated knockdown of tsRNAs (Fig. S1D) laying credence to the idea that tsGlnCTG specifically targets transcripts other than neural lineages during RA-induced differentiation. Taken together, these data suggest that upon RA treatment tsGlnCTG functions by sequestering distinct subsets of mRNAs important for pluripotency and non-neuronal lineage specification allowing the stem cells to continue along the path of neural differentiation.

**Figure 3.**
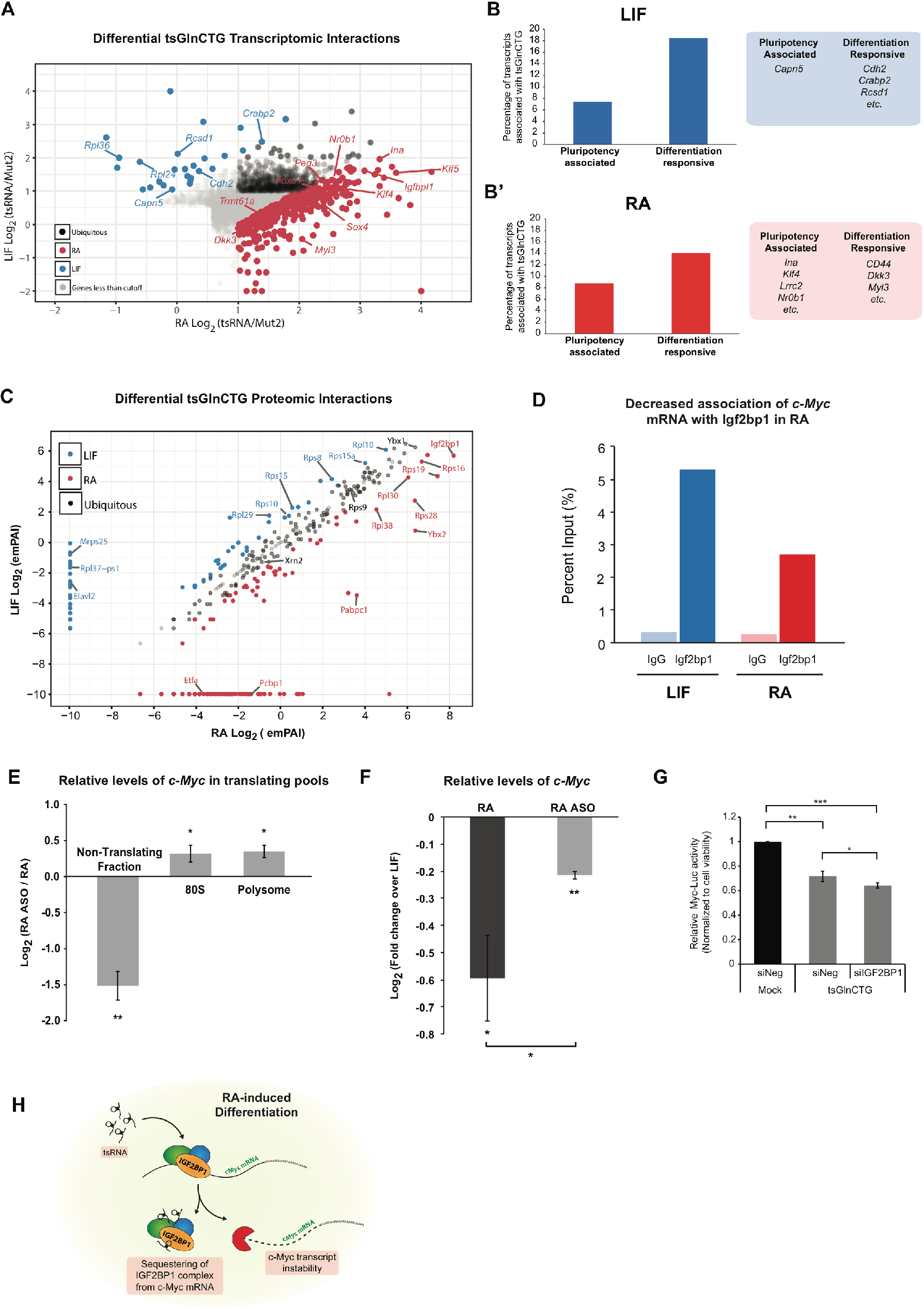
Identification and functional characterization of tsRNA interactome in mESCs cultured with LIF or RA. **(A)** Scatter plot showing Log_2_ fold-change enrichment of mRNAs associated with tsGlnCTG in LIF-versus RA-treated mESCs. **(B and B’)** Bar plots and representative gene lists depicting association of tsGlnCTG with ‘pluripotency-associated’ (B) and ‘differentiation-responsive’ (B’) genes in LIF- or RA-treated mESCs. **(C)** Scatter plot showing Log_2_ fold-change enrichment of protein interactome between LIF- and RA-treated mESCs. All peptides identified by LC-MS/MS had < 1% FDR (Fig. S3A). LC-MS/MS was conducted twice (Fig. S3B, correlation analysis). **(D)** Quantification of Igf2bp1 bound *c-Myc* mRNA enrichment between LIF- versus RA-treated mESCs (Refer Fig. S3G’ for duplicate data) **(E)** Relative enrichment of *c-Myc* mRNA in translating (80S and Polysome) and non-translating pools (mRNPs) fractionated from polysome profiling between ASO-treated and Mock-treated RA-induced differentiating mESCs. (Error bars represent SD, significance was calculated using one-tailed unpaired t-test) **(F)** Relative levels of *c-Myc* mRNA in ASO-treated and Mock-treated RA-induced differentiating mESCs in comparison to LIF (Error bars represent SD, significance was calculated using one-tailed unpaired t-test) **(G)** Epistatic analysis of tsRNAs and IGF2BP1 in regulating Myc transcriptional reporter activity. *, *P* value < 0.05, **, *P* value < 0.01, ***, *P* value < 0.001, *n.s.*, not significant.

The association of tsRNAs to mRNAs could be either direct (sequence complementarity), or indirect (through associating with certain RNA binding proteins). Bioinformatics analysis failed to identify any significant complementarity between tsGlnCTG and the mRNAs (data not shown) suggesting that the interaction could be through tsRNA-associated proteins. We therefore conducted pull-down assays followed by mass spectrometry using the tsGlnCTG mimic in mESCs treated with either LIF or RA to identify differentially interacting proteins (Fig. S3A). Proteomic analysis revealed that tsGlnCTG interacted with several ribosomal proteins and RNA-binding proteins (RBPs) in stem versus differentiating state (Fig. 3C and Table S5). Interestingly, we observed distinct subsets of ribosomal proteins that interact with tsGlnCTG in stem versus differentiating states. While it has been well established that ribosomal heterogeneity plays an important role in regulating translation of specific transcripts (Xue and Barna, 2012), our data invokes the intriguing possibility that tsRNAs impose target selection by virtue of associating with specific ribosomal/RNA binding proteins in different cell states.

### tsRNAs regulate c-Myc transcript levels through association with Igf2bp1

In RA-treated mESC we observed a strong association of tsGlnCTG with the RNA binding protein Igf2bp1/Imp1 (Fig. 3C and S3B), a known factor required for stem cell maintenance and function (Conway et al., 2016, Nishino et al., 2013). Although the levels of Igf2bp1 over 48hrs of RA-induced differentiation remains constant (Fig. S3E and S3E’), RNAi-mediated knockdown of Igf2bp1 led to a loss of pluripotency. This was evident from decreased expression of pluripotency genes upon Igf2bp1 knockdown (Fig. S3F and S3F’). Igf2bp1, is also known to prevent endonucleolytic cleavage of c-Myc mRNA by binding to Coding Region instability Determinant (CRD), thus facilitating efficient c-Myc mRNA association with the translating pool (Weidensdorfer et al., 2009). c-Myc is a regulator of cell states; its overexpression induces pluripotency (Okita et al., 2008), and its loss results in entry into a diapause state (Scognamiglio et al., 2016). We hypothesized that the change in interactions of Igf2bp1 with c-Myc could be crucial for maintaining pluripotency or differentiation. Therefore, we investigated whether the increased association of tsRNAs with Igf2bp1 perturbs its binding to c-Myc mRNA facilitating the transition from pluripotency to differentiation.

We performed RNA immunoprecipitation (RIP) assays to study the interaction of Igf2bp1 with tsRNAs and the c-Myc transcript. RIP followed by small RNA sequencing from RA-treated mESCs revealed associations of Igf2bp1 with several tsRNA including tsGlnCTG, validating our tsGlnCTG–protein interactome data (Table S6 and Fig. S3G). Additionally, by performing real-time PCR analysis of the Igf2bp1 RIP, we observed a decreased association of Igf2bp1 with *c-Myc* mRNA during differentiation compared to LIF condition (Fig. 3D and Fig. S3G’). To investigate the crucial role of tsRNAs in regulation of c-Myc-Igf2bp1 association, we blocked tsRNAs using ASOs during RA-induced differentiation. Notably, polysome profiling revealed decreased levels of *c-Myc* mRNA association with actively translating pools (Polysomes and 80S) in RA treated cells compared to LIF-treated ESCs (Fig. S3H). However, when tsRNAs were blocked using ASOs in RA-treated ESCs, we observed increased *c-Myc* association with translating pools with a simultaneous decrease in the nontranslating pool (Fig. 3E). This was further validated by the data that showed an overall increased levels/stability of *c-Myc* mRNA in RA-ASO compared to RA-treated condition (Fig. 3F). To further elucidate the antagonistic role of tsRNAs in blocking the interaction between Igf2bp1 and *c-Myc*, we used HCT116 Myc-Luc reporter cell lines to measure the luciferase activity driven by a c-Myc regulated promoter. The extent of luciferase activity is a direct measure of levels of c-Myc protein. We transfected tsGlnCTG into the HCT116-Myc-Luc reporter cell line in the background of IGF2BP1 knockdown (siRNA). Evidently the overexpression of tsGlnCTG was sufficient to inhibit luciferase activity by ∼30% which was further reduced by an additional 10% upon IGF2BP11 knockdown (Fig. 3G). Taken together, these data suggest a model for tsRNA function in modulating the binding of Igf2bp1 to *c-Myc* mRNA which is crucial for the stability and translation of *c-Myc* transcript (Fig. 3H).

### tsRNA expression is a global hallmark of stem/non-stem states

To investigate tsRNAs as global regulators of cell-state transitions, we performed small RNA sequencing on a variety of well-defined models for stem/transformed versus differentiated/normal states. The stem/differentiation pattern of tsRNA expression was observed in a variety of contrasting cell states across heterologous model systems. We looked for tsRNAs *in vivo*, in sorted stem and differentiating cell populations of murine skin. In agreement with the tsRNA-differentiation bias observed in mESCs, CD34^−^α6^+^ (non-stem) cells showed a 11% increase in a discreet 32-mer tsRNA population, compared to the CD34^+^α6^+^ multipotent stem compartment (Fig. 4A, Tables S1 and S2). Remarkably, the tsRNA levels were upregulated in the differentiating basal cells compared to the bulge stem cells, but were downregulated in the terminally differentiated suprabasal keratinocytes (Fig. S4A), parallel to expression observed in mESC differentiation (Fig. S2A). Next, we examined human mammary epithelial cells (HMECs) at different stages of oncogenic transformation (Cheong et al., 2011; Voorhoeve and Agami, 2003) (Fig. S4B and B’, validation of cell lines). As HMECs transform, they acquire stem cell-like morphologies (Fig. 4B), display expression of stem markers (Fig. S4C, gain of *CD44* expression and loss of *CD24)* and enhanced EMT signatures (Fig. S4D). We detected > 50% decrease in levels of 30-35 nt RNAs (by Bioanalyzer electrophoresis) in the transformed, poorly differentiated cancer “stem” cells compared with the “normal” cell state (Fig. 4B’). Notably, the decrease in small RNA levels appeared to be an early cell transformation event. Additionally, a differential tsRNA profile in CD44^+^CD24^−^ (stem) and CD24^+^ (non-stem) cells was observed in two breast cancer cell lines by small RNA sequencing. Specifically, the CD24^+^ cells in both MDA-MB-231 and HS578T showed a 15% and 9.3% increase respectively in a specific 33-mer tsRNA population compared to the CD44^+^CD24^−^ cells (Figs. 4C and 4D).

**Figure 4.**
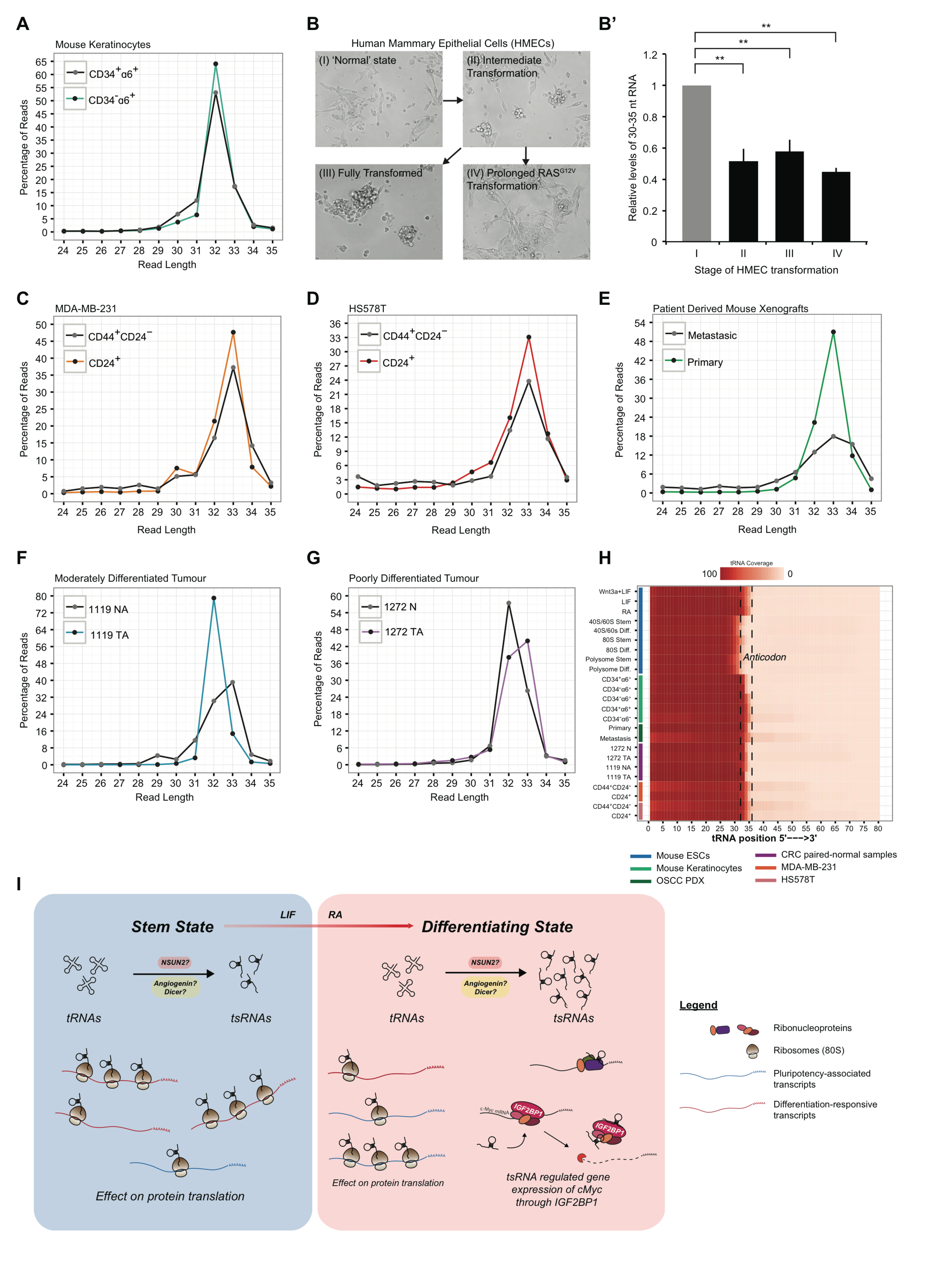
Global association of tsRNA expression with cell state switches. **(A)** tsRNA profile of bulge stem cell (CD34^+^α6^+^) or basal cell (CD34^−^α6^+^) populations isolated from adult murine skin (Plot S1). **(B)** Images of HMECs (Panels I-IV) engineered to represent stages of oncogenic transformation: I: parental HMECs immortalized with hTERT; II: immortalized HMECs with tamoxifen-inducible oncogenic HRAS (hTERT/sT/HRAS^G12V^:ER/EV); III: HMEC as described in II with additional p53 knockdown (hTERT/sT/HRAS^G12V^:ER/shP53); HMECs I-III were treated with 130 nM tamoxifen (5 days). IV: HMEC IIs treated with tamoxifen (10 days). (B’) Quantification of 30-35 nt RNA levels in HMEC I-IV. **(C and D)** tsRNA profiles of stem (CD44^+^CD24^−^) versus differentiated (CD24^+^) populations sorted from breast cancer cell lines; MDA-MB-231 (C) and HS578T (D) (Plots S2 and S3). **(E)** tsRNA profiles of Patient-Derived mouse Xenograft lines from primary or metastatic OSCCs. **(F and G)** tsRNA profiles of paired-normal CRC patient resected tumours with either a moderately differentiated (F) or poorly differentiated (G) tumor. N: Normal; NA: Normal Anterior; TA: Tumor Anterior. **(H)** Heat map of tsRNA reads mapped to parent tRNA residues across various cellular systems sequenced in this study. Majority of the tsRNAs identified were processed specifically from the 5’-half of the mature tRNA **(I)** Proposed mechanistic model for tsRNA cellular and molecular functions. **, *P* value < 0.01.

To extend the scope of our investigations beyond animal models and established cell lines, we probed for differential tsRNA profiles in clinically relevant models. In a pair of patient-derived xenograft (PDX) lines originating from synchronous primary and metastatic oral squamous cell carcinoma (OSCC) (*Chia et al., 2017, in press*), we found a 33% decrease in the 33-mer tsRNA population in the metastatic cells compared to primary/parental tumor cells (Fig. 4E). This is consistent with previous findings by Goodarzi *et al.* that breast cancer cells with enhanced metastatic properties express lower levels of tsRNAs (Goodarzi et al., 2015). Detecting levels of tsRNAs may be a clinically relevant diagnostic tool, to complement histological grading of tumors. We examined two contrasting sets of paired resected tumors from colorectal cancer (CRC) patients 1119 and 1272. Patient 1119 had a moderately differentiated tumor (1119TA), whereas sample 1272TA was histologically classified as poorly differentiated. Further molecular validation indicates that 1119TA express lower levels of CRC stem markers compared to paired normal, and markedly higher levels of pro-differentiation markers (Fig. S4E). The opposite was observed in 1272TA (Fig. S4F). Small RNA sequencing revealed that 1119 tumor expressed 48% more, while 1272 tumor had a 19.3% reduction of 32-mer tsRNAs compared to their respective paired normal samples (Figs. 4F and 4G). Collectively, tsRNAs are hallmarks of cellular and cancer differentiation.

## Discussion

Cell state transitions are fundamental to normal development. The process of stem cell differentiation into lineage-specific progenitors involves changes at both transcriptional and translational levels. In this present study, we have identified a role for tsRNAs in modulating translation, which is critical to the differentiation of mESCs along the neuro-ectodermal lineage upon RA treatment. In our study, we identify a pool of tsRNAs that modulate differentiation of mESCs. Although several studies have identified diverse tRNA-derived small RNA species in different biological contexts (Goodarzi H, et al. 2015; Lee SR, et al. 2005; Chen et al., 2016; Sharma et al., 2016), our study elucidates the function of tsRNAs derived from 5’ half of tRNAs in stem cell differentiation. We investigated the nature of these tsRNAs across various cell-state-transition contexts and observed that the majority of tsRNAs specifically and consistently matched to the 5’ half of tRNAs (Fig. 4H).

We show that the tsRNAs enriched during differentiation inhibit translation in agreement with the previous reports (Fig. 2D) (Ivanov, P, et al, 2011; Sobala A et al, 2013). We also observed a significant enrichment of 5’ tsRNAs in monosome and polysome fractions during differentiation (Fig. 2E), suggesting that tsRNAs may regulate translation at multiple steps in addition to their reported role in repressing translational initiation (Ivanov P, et al. 2011). Furthermore, biochemical assays revealed specificity in the interaction of tsRNAs with the transcripts that are known to regulate stemness and differentiation. Notably, tsRNA association with “differentiation-responsive” transcripts in the stem state suggests a function in translational repression of transcripts that are critical for maintenance of stemness (Fig. 3B). However, during RA-induced differentiation, tsRNAs interact with both “pluripotency-associated” and “differentiation-responsive” transcripts (Fig. 3B’). Although the association of tsRNAs with “differentiation-responsive” transcripts during differentiation is counter intuitive, GO annotation revealed that the transcripts were enriched for several lineages apart from those promoting or associated with the neuroectoderm (Table S4). Based on our findings, we propose a role for tsRNAs in translational suppression of transcripts critical for the maintenance of stem cell state or lineage specification during differentiation.

As evident from the work by Goodarzi et al., the tsRNA-protein interaction is critical for target selection. Our study showed tsRNAs largely interact with proteins such as RNA-binding proteins and ribosomal proteins involved in the regulation of mRNA stability and translational repression (Fig. 3C and Table S5). The associations of tsRNAs with different sets of ribosomal proteins between stem versus differentiating cells invoke two exciting possibilities with respect to tsRNA function. First, tsRNAs may associate with specific pools of ribosomes (and hence ribosomal proteins) that are involved in the translation of specific targets. Second, the spatial positioning of tsRNAs on the ribosome, evident from its interaction with both large and small subunit, suggests different modes of translational regulation. Detailed structural studies may be required to better understand the tsRNA associations with ribosomal proteins.

Here, we focused on the importance of one such interaction; specifically, between the tsRNA, tsGlnCTG, and Igf2bp1, and its functional consequences in the regulation of c-Myc, an important pluripotency factor. Igf2bp1 is an RNA-binding protein implicated in stem cell maintenance and carcinogenesis that regulates the stability and translation of several mRNAs (Conway et al., 2016, Wang, G et al 2016). IGF2BP1 was shown to stabilize the transcript, thereby promoting efficient translation (Weidensdorfer et al., 2009). Since our tsGlnCTG-protein interactome revealed binding of tsRNA to Igf2bp1 in RA conditions (Fig. 3C), we hypothesized that tsRNAs could potentially regulate *c-Myc* through its association with Igf2bp1. In agreement with our hypothesis, we found decreased association of the c-Myc transcript with Igf2bp1 in RA-induced differentiating cells (Fig. 3D), suggesting that the interaction of tsRNAs (such as tsGlnCTG) with Igf2bp1 could potentially destabilize the *Igf2bp1-c-Myc* interaction. Loss of function studies using ASOs resulted in increased c-Myc mRNA stability and its association with the translating pools, validating our hypothesis that tsRNAs dictate the fate of c-Myc mRNA through their interaction with Igf2bp1 (Fig. 3G). This is further supported by gain-of-function studies using tsRNA mimics, where we observed decreased luciferase activity under the influence of the promoter driven by c-Myc. Altogether our results suggest that tsRNAs play a critical role in regulating c-Myc mRNA stability through its interactions with Igf2bp1 to modulate stem cell differentiation.

It has been shown that Igf2bp1 binds to different regions on the mRNA; 5’UTR, CDS or 3’UTR, essential for determining the fate of the mRNA (Conway et al., 2016). It is interesting to note that tsRNAs interact with several transcripts, some of which could be targeted through Igf2bp1 that requires further characterization. Previous reports have shown tsRNA interaction with YBX1 is critical regulating transcript stability and repression of translational initiation (Goodarzi H et al., 2016; Ivanov, P et al., 2011). Though in the current study we found tsRNA interaction with YBX1, we did not observe differential interaction with YBX1 between stem versus differentiating states (Fig. 3C and Table S5). This suggests that tsRNA-YBX1 interaction may be involved in regulating transcripts essential for both self-renewal and differentiation. Probing tsRNA interaction with other RNA-binding proteins will likely provide a comprehensive understanding of tsRNA function during stem cell differentiation. Taken together our data suggests a bimodal role for tsRNAs in stem versus differentiation state; wherein, tsRNA in the stem state potentially regulate the ‘differentiation responsive’ transcripts and upon induction to differentiate, tsRNAs regulate a different set of transcripts critical for lineage commitment thereby facilitating differentiation (Fig. 4I).

Finally, while earlier studies have identified tsRNAs in cancer cell lines (Goodarzi, H, et al., 2016), in the present study we probed tsRNA expression in patient-derived paired-normal versus tumor tissue. Comparison of tsRNA profiles between normal and tumor tissue of distinct clinical grades revealed the preferential enrichment of a specific 33 nt pool of tsRNAs in the more differentiated tumors (Fig. 4F and 4G). Similar enrichment of 33 nt tsRNAs was also observed in primary tumors compared to metastatic tumors derived from PDX models (Fig. 4E). These observations suggest processing of tRNAs at a specific position to generate 33 nt tsRNAs during tumor progression that involves cell-state transitions. This enrichment of 33 nt species of tsRNA could be exploited as potential biomarker for cancer progression. Further, understanding the functional relevance of 33 nt tsRNA species in the tumor progression/differentiation may facilitate the development of RNA-based anti-cancer therapeutics.

In summary, our data provides evidence that tsRNAs interact with specific sets of mRNAs critical for cell state transitions. Further we also provide a new translational regulation model based on tsRNA-Igf2bp1 interaction, which is essential for cellular differentiation.

## Methods

### Cell culture

E14 mouse embryonic stem cells (mESCs) were cultured in high glucose DMEM media, supplemented with 15% ES grade fetal bovine serum (FBS) (Biowest S182S-500, Batch S10397S182), sodium pyruvate, gentamycin (Gibco 15710-064), non-essential amino acids (NEAA) (Gibco 11140-050), glutamax (Gibco 35050-061), ß-Mercaptoethanol (Gibco 21985023) and 1000 U/mL LIF (ESGRO 1106). The mESCs were grown on 0.1% gelatin coated plastic and passaged every 2 days, with daily media changes, and for a maximum of 12 passages. mESCs were differentiated with 0.5 µM RA for 48 hrs, unless specified otherwise.

HCT116 Myc-Luc reporter cells were purchased from Amsbio, BPS Bioscience (#60520). MDA-MB-231 and HS578T breast cancer cell lines were gifts from Wai Leong TAM. The above were cultured in high glucose DMEM with sodium pyruvate, supplemented with 10% FBS (Biowest S181B-500), 1% penicillin/streptomycin (Gibco 15140-122), and 1% glutamax (Gibco 35050-061). The HCT116 Myc-Luc reporter line is additionally cultured under selection with 500 μg/ml Geneticin. Human Mammary Epithelial Cells (HMECs) 1-3 were a gift from Mathijs Voorhoeve. The HMECs were maintained in MEGM™ Mammary Epithelial Cell Growth Medium (Lonza MEGM BulletKit (CC-3151 & CC-4136)).

### RNA isolation

Total RNA from cells, including small RNAs were isolated through phase separation with TriZol according to manufacturer’s recommendations, followed by capture with the miRNeasy kit (Qiagen 217004). Frozen patient tumors were homogenized in gentleMACS™ M Tubes (MiltenyiBiotec 130-093-236) on a gentleMACS Dissociator (130-093-235), using program RNA_02. Total RNA from the dissociated tumors cells was then isolated with TriZol and the miRNeasy kit.

### Quantitative real-time PCR (qRT-PCR)

Total cellular RNA was extracted as described above. Atleast two biological replicates were used per sample. Reverse transcription (RT) was conducted with the SuperScript II Kit (Invitrogen) according to the manufacturer’s protocol. qPCR utilizing the KAPA SYBR FAST qPCR mix (KAPA Biosystems) was conducted with the QuantStudio® 7 Flex Real-Time PCR System (ThermoFisher Scientific). Primers for qRT-PCR reactions are listed in Table S7.

### Small RNA Library Preparation

All libraries generated for small RNA sequencing was prepared using TruSeq Small RNA Prep Kit (RS200012, RS200024, RS200036, RS200048, Illumina) using the manufacturers protocol. The libraries were multiplexed and sequenced on Illumina NextSeq 500 platform.

### Identification and characterization of tsRNAs

We used mm10 (Chinwalla AT et al., 2002) and hg19 (Lander ES et al., 2001) as the reference genomes (http://genome.ucsc.edu/) for mouse and human samples respectively. From the sequencing reads, we trimmed TruSeq small RNA adapters using customized perl script and cutadapt program (Martin M., 2011). The adapter trimmed reads were aligned to rRNA and unaligned reads were taken for further analysis. We then segregated reads that are 18-35 nucleotides and mapped to Genome and tRNA (Chan PP et al., 2008) databases using bowtie v1.0.0 (Langmead B et al., 2009). We used 2 mismatches as constant parameter for all the alignments done in this study, as we did not observe much deviation in our results with varying mismatches ranging from 0-2 (Table S1). Reads, which are 25-35 nucleotides in length and specifically maps only to tRNA database, were considered as tsRNAs. We did not consider 18-24mer reads as tsRNAs because only negligible amount mapped to tRNA database. We calculated per base tRNA coverage using the following formula: *Per base coverage* = (number of reads aligned to particular base of tRNA/total number of reads mapped to all the tRNA in that sample).

We used VARNAv3-93 (Darty K et al., 2009) for obtaining secondary structures of tRNA and we loaded the obtained coverage values to get tRNA heatmap. We used DESeq (Anders S et al., 2010) for normalization and identification of differentially expressed tsRNAs (adj. *P*-value < 0.05). All the statistical tests were done in R (R Core Team). We used customized perl script for all the analysis used in this study. We used R ggplot2 (Wickham H, 2010) library for plotting. The raw numbers and mapping percentage for all the sequenced samples are given in Table S1.

### Small RNA qPCR

Total RNAs isolated from mESCs (LIF- or RA-treated). Atleast two biological replicates were used per sample. For each sample, 1 µg of RNA was then reverse transcribed with Superscript II RT (Invitrogen) using tsRNA specific stem-loop RT primers (Sigma) (Table S7). Reverse transcribed cDNA was amplified using Maxima SYBR Green/ROX qPCR Master mix (K0222, Fermentas) using tsRNA specific forward primer and universal reverse primer against the stem-loop RT primer (Table S7). Sno-202 was used as an endogenous control to normalize differences. The samples were amplified and analyzed on Applied Biosystems 7900HT qPCR machine.

### RNA oligonucleotides

All RNA oligonucleotides were synthesized and purchased from Integrated DNA Technologies. RNA oligonucleotides against sequences of candidate tsRNAs are 2’OMe modified. Chimeric Antisense Oligonucleotides (ASOs) are complementary phosphorothioate modified DNA sequences flanked by 2’OMe modified RNA. Pool of 5 ASOs against tsGlyGCC, tsGlyCCC, tsGlnCTG, tsValCAC and tsArgTCT were used to investigate the effects of tsRNA knockdown. 3’-Biotin with TEG linkers were tagged to tsGlnCTG or a nonsense scrambled control (Mut2). Sequences of oligonucleotides used are found in Table S7.

### Transfections of mESCs

3 x 10^5^ or 6 x 10^5^ mESCs per well (6-well plate) were plated for forward and reverse transfections respectively. mESCs were transfected with Lipofectamine 2000 (ThermoFisher Scientific) according to manufacturers’ recommendations. mESCs transfection media consists of 1:1 parts of 2x mESC media supplemented with LIF, and OptiMEM Reduced Serum Media (Gibco 31985070). Transfection media was changed to mESC media without antibiotics 5-6 hrs or 16-18 hrs post transfections for forward and reverse transfections respectively.

Alternatively, for large scale experiments such as polysome profiling followed by QPCR, cells were transfected in Neon Transfection system (Thermo Fischer, MPK10096) at 1100 volts for 3 pulses. 1 uM concentrations of each tsRNA antisense (ASO) was used for transfection.

### Alkaline Phosphatase activity assay

Alkaline Phosphatase activity of mESCs was measured using the Alkaline Phosphatase Assay Kit (Colorimetric) (Abcam ab83369), in accordance to product recommended protocol. The mESCs were grown and transfected in 48-well TC plates. Optical densities at 405 nm were read with the Tecan Infinite M1000 PRO Multimode Microplate Reader in 96-well clear flat bottom TC plates.

### Microfluidic RNA gel electrophoresis

Equal amounts (not exceeding 100 ng) of total RNA per sample was loaded in each small RNA chip well (Agilent Small RNA kit, 5067-1548), in accordance to product protocols, and analyzed with the Agilent 2100 Bioanalyzer. Nucleotide size regions of interest were gated and subsequent quantitation of RNA abundance of specified regions were obtained using the Bioanalyzer Expert software.

### Polysome profiling

Mouse ESCs (4.86 x 10^6^) were seeded in each 15 cm dishes per culture condition. Translation was arrested by incubating cells with 100 μg/ml cycloheximide (CHX, Sigma # C4859) for 10 min at 37°C. After washing cells on ice with PBS supplemented with 100 μg/ml CHX, cells were lysed in buffer (10 mM Tris-HCl (pH 7.4), 5 mM MgCfe, 100 mM KCl, 1% Triton X-100, 2 mM DTT, 500 U/ml RNas Inhibitor, 100 μg/ml CHX and Protease inhibitors). Cell lysates were then sheared gently four times with a 26-gauge needle. Lysates were then collected after centrifugation at 1300 x g for 10 min. Lysates were layered onto 10–50% sucrose gradients and centrifuged in an SW-41Ti rotor at 36,000 r.p.m. for 2 hrs. Gradients were fractionated using a BioComp Gradient Station fractionator, and absorbance at 254 nm was monitored to obtain the polysome profile. Polysome:monosome (P:M) ratios were derived by integrating the area under the respective peaks.

### Click-iT labelling and detection

Three hours before the indicated time points, cells were washed with warm PBS and incubated in methionine-free DMEM media with 15% dialyzed FBS for 1 hr, at 37 °C to deplete methionine reserves. Click-iT AHA (L-Azidohomoalanine, C10102, Thermo Fisher Scientific) was then added at a final concentration of 50 µM, for 3 hrs at 37°C, for metabolic labelling of newly synthesized proteins. At 1 hr past the indicated time points, cells were harvested by scraping in PBS. Cells were then lysed in lysis buffer (see above polysome profiling methods section). The lysate was homogenized six times with a 26-gauge needle at 4 °C and centrifuged at 1,300g for 8 min. Proteins were then extracted from the supernatant by methanol/chloroform precipitation. AHA-labelled proteins were detected by Click chemistry with tetramethylrhodamine alkyne (TAMRA, T10183, Thermo Fisher Scientific) using the Click-iT protein reaction buffer kit (C10276, Thermo Fisher Scientific), according to manufacturers’ instructions. Proteins were then separated by SDS-polyacrylamide gel electrophoresis (SDS-PAGE). The TAMRA signal was visualized using the ChemiDoc MP Imaging System (Bio-Rad), with the setting to detect Alexa Fluor 546. The same gel was then stained with Coomassie to visualize total proteins. Relative protein synthesis levels were derived by normalizing TAMRA signal to the Coomassie signal.

### In-vitro translation assay

For *in vitro* transcription, EGFP was amplified from pEGFPN3 construct using a T7-tagged Forward primer and a reverse primer (Table S7). Capped-polyadenylated GFP mRNA was synthesized using mMESSAGE mMACHINE T7 Ultra Kit (AM1345, Ambion) following manufacturer’s protocol. 100-200 ng of the capped-polyadenylated GFP mRNA was used in Retic Lysate system (AM1200, Ambion) and incubated at 30 °C for 90 mins. Scramble or tsGlnCTG (Table S7) was added to the lysate at varying concentrations. Emission of GFP at 507 nm was measured using a spectrofluorometer (Fluorolog3, Horiba Scientific).

### Small RNA Sequencing from Polysomes

Prior to polysome profiling, cells were translationally arrested by treating with CHX for 30 minutes. Lysates made from vairous states of differentiation (*Stem: Wnt3a+LIF- and LIF-treated; Differentiating: RA- and Wnt3a+RA-treated*) were ultra-centrifuged on a 15-45% sucrose gradient at 39000 r.p.m. to separate the ribosomal fractions. The individual subunits (40S, 60S), monosomes (80S) and polysomes were fractionated based on the profile generated by the absorbance of the gradient at 254 nm. The RNA was precipitated from the sucrose gradient collected by ethanol precipitation. The pellets were dissolved in TriZol and RNA was isolated from the pellets. Small RNA libraries were prepared from isolated RNA using TruSeq Small RNA Prep Kit (RS200012, Illumina). For statistics, samples treated with Wnt3a+LIF and LIF were grouped as proxies for stem, and RA- and Wnt3a+RA-treated (absence of LIF) as proxies for differentiating state. The data was analysed as described in Methods Section “Identification and characterization of tsRNAs”. We used Wilcox’s test (R Core Team) to calculate significance. The change in the tsRNA profiles were calculated using change point analysis (R package: changepoint) (Killick R and Eckley I, 2014).

### Electron microscopy

In vitro translation lysates were taken for electron microscopy sample processing after 90 mins of incubation at 30 °C. The lysates were coated onto Holey-Carbon grids for 5 mins and washed thrice with water. Grids were stained with 2% Uranyl acetate for 30 seconds and washed thrice with water and allowed to dry. The grids were imaged using Technai G2 Spirit Bio-TWIN transmission electron microscope (FEI).

### Biotinylated RNA mimics pulldown

Pulldowns with 3’-Biotinylated RNA mimics were conducted with a protocol adapted from (Tsai MC et al., 2010). 3 μg of folded biotinylated tsGlnCTG or a scrambled sequence (Mut2), in 100uL of RNA structure buffer (10 mM Tris pH 7.0, 0.1 M KCl, 10 mM MgCl_2_), were incubated with 1 mg of pre-cleared cell extracts in cell lysis buffer (20 mM Tris-HCl pH 7.5, 150 mM NaCl, 1.8 mM MgCl_2_, 0.5% NP40, Protease inhibitor cocktail for mammalian cells (no EDTA), 1 mM DTT, 80 U/mL RNase inhibitor). Proteins were eluted with SDS buffer from Streptavidin agarose beads, and analysed on PAGE gel electrophoresis. RNA was extracted from Dynabeads M-280 Streptavidin™ (11206D, Invitrogen) beads using TriZol (Invitrogen). Isolated RNA was analyzed on microfluidic RNA gel electrophoresis with the Agilent 2100 Bioanalyzer before RNA sequencing.

### tsRNA mediated mRNA pulldown library and analysis

The paired-end sequencing reads were adapter trimmed using Trimmomatic and aligned to mouse (mm10) genome using Tophat v2.0.9 (-p 10 -o ./ -i 50 -I 250000) (Trapnell C et al., 2009). Around, 92-95% of reads mapped back to reference genome. We used HTSeq-count v0.6.0 (Anders S et al., 2015) (-f bam –r name –s no –m union -o) to obtain raw read numbers for all the Ref Seq transcripts. Gene body coverage analysis, for all samples, showed reads covering the entire transcript and not from one specific region. The count data was normalized using DESeq. We used a cut-off of DESeq normalized value > 200 in at least one of the four samples (Mut2 and tsRNA pulldown in LIF and RA). We further selected transcripts that showed tsRNA/Mut2 fold change ratio to be more than 2 fold. To identify transcripts that significantly associate to tsRNA in comparison to Mut2, we used Fisher exact test (Fay MP, 2009) (fisher.test (x,alternative=“two.sided”,simulate.*P*.value = FALSE,B = 2000)). *P*-value obtained from Fisher exact test was corrected using *Bonferroni* method (stats package in R) (R Core Team). Transcripts with adjusted *P*-value < 0.05 were considered as significant. Either in LIF or RA, 3664 transcripts showed significant association in tsRNA pulldown compared to their respective control (Mut2) (Table S3).

Further, to identify cell-state specific tsRNA associated transcripts we pruned this list based on tsRNA_RA/tsRNA_LIF fold-change values. A cut-off of two fold was used to allocate transcripts as either RA enriched (420 transcripts) or LIF enriched (28 transcripts). We mined publicly available transcriptome data (Chatterjee SS et al., 2015) and classified transcripts that were more than two fold up-regulated after RA treatment as “differentiation responsive genes” and more than two-fold downregulated as “pluripotency associated genes”. Genes that showed tsRNA/Mut2 was >2 fold in both the LIF and RA conditions and similar enrichments in both the conditions were categorized as ‘Ubiquitous’. Subsequently we correlated our tsRNA pull-down data with these two classes of gene sets to pinpoint molecular players that tsRNA might most possibly be regulating and interacting. The GO analyses for different catogories were done using g:profiler (http://biit.cs.ut.ee/gprofiler/) (Reimand J et al., 2016).

### Protein Elution and In-Gel Tryptic Digestion

The beads used to pull down the RNA-protein complex were boiled in gel loading buffers. The eluted samples were run on a SDS-PAGE gel. Gel slices were cut into small pieces and transferred to Eppendorf tubes. They were washed for several time with Milli-Q water, and then followed with 50% ACN/50% 25mM NH_4_HCO_3_ via vigorous vortexing for 30 min. The gel pieces were then dehydrated with 100% ACN. They were then reduced with 10 mM DTT (dithiothreitol) at 56 °C for 1 hr and alkylated with 55 mM IAA (iodoacetamide) for 45 min in the dark followed by successive washes with 25 mM NH_4_HCO_3_ and 50% ACN/50% 25 mM NH_4_HCO_3_ with vigorous vortexing for 30 min. The gel pieces were dehydrated again with 100% ACN. Trypsin (V5111, Promega, Madison, WI) was added in the ratio of 1:50. After the trypsin solution was completely absorbed by gel particles, 25 mM NH4HCO3 was added to completely cover the particles. They were then incubated at 37 °C overnight. Peptides were extracted from gel particles with 50% ACN containing 0.1% TFA under sonication for 30 minutes twice. The combined extracts were dried in vacuum and stored at −20 °C before LC-MS/MS analysis.

### LC-MS/MS

LC-MS/MS was done as previously described (Cheow ESH et al., 2016). Tryptic peptides were dissolved in 0.1% formic acid (FA) in 2% Acetonitrile (ACN). They were then analyzed on a Dionex Ultimate 3000 RSLC nano system coupled to a Q Exactive tandem mass spectrometer (Thermo Fisher, CA). Each peptide sample was injected into an Acclaim peptide trap column (Thermo Fisher, CA) via the auto-sampler of the Dionex RSLC nano system. Peptides eluted from the peptide trap were separated in a Dionex EASY-spray column (PepMap^®^ C18, 3um, 100A, 75 µm x 15 cm) (Thermo Fisher, CA) at 35 °C. Mobile phase A (0.1% FA in H_2_O) and mobile phase B (0.1% FA in 100% ACN) were used to establish a 60 min gradient at a flow rate of 300 nl/min. Peptides were then analyzed on Q Exactive with an EASY nanospray source (Thermo Fisher, MA) at an electrospray potential of 1.5 kV. A full MS scan (350-1600 m/z range) was acquired at a resolution of 70,000 at m/z 200 and a maximum ion accumulation time of 100 msec. Dynamic exclusion was set as 30 s. Resolution for HCD spectra was set to 35,000 at m/z 200. The AGC setting of full MS scan and MS^2^ were set as 1E6 and 2E5, respectively. The 10 most intense ions above a 1000 counts threshold were selected for HCD fragmentation with a maximum ion accumulation time of 120 msec. Isolation width of 2 Th was used for MS^2^. Single and unassigned charged ions were excluded from MS/MS. For HCD, normalized collision energy was set to 28%. The underfill ratio was defined as 0.1%.

### LC-MS/MS Data Analysis

The raw data were processed as previously described (Adav SS et al., 2016). Briefly, raw data files were converted to the mascot generic file format using Proteome Discoverer version 1.4 (Thermo Electron, Bremen, Germany) with the MS2 spectrum processor for de-isotoping the MS/MS spectra. The concatenated target-decoy Uniprot (Apweiler R et al., 2004) human database (sequence 88 473, downloaded on 29 November 2013) was used for MS/MS spectra searches. The database search was performed using an in-house Mascot server (version 2.4.1, Matrix Science, Boston, MA) with MS tolerance of 10 ppm and MS/MS tolerance of 0.02 Da. Two missed trypsin cleavage sites per peptide were tolerated. Carbamidomethylation (C) was set as a fixed modification, while oxidation (M) and deamidation (N and Q) were variable modifications. Label-free quantitation of proteins were based on emPAI values of each identified proteins reported by Mascot. The relative protein quantities between samples were then calculated from the emPAI values of the protein in the samples.

### Analysis of tsRNA proteomic associations

Non-detectable emPAI (protein abundance index) values (-1) were replaced with lowest detected values (0.01). Beads emPAI values were then subtracted from tsGlnCTG emPAI values. Candidates that display higher emPAI scores in bead controls versus respective conditions were disregarded. Specific functional enrichment of tsGlnCTG associated proteins showing more than a 2-fold change in emPAI scores were investigated for enrichment of Molecular Signature Gene sets (MSigDB, v5.1) (Subramanian A et al., 2005) using Fisher’s Exact Test, followed by multiple testing correction using False Discovery Rate (FDR) estimation (Benjamini Y and Yekutieli D, 2001). Significant gene sets were determined at FDR adjusted P-value < 0.05 (Table S5).

### Western blotting

After PAGE gel electrophoresis and PVDF membrane transfer using standard laboratory protocols, antibodies against IMP1/IGF2BP1 (Cell Signaling Technologies D33A2, #8482S) and ß-Actin (Abcam 8226, ab8226) were used to probe for target proteins. IR Dye 680RD Goat anti-Rabbit (LiCor 926-68071, Lot# C31205-05) and IR Dye 800CW Goat anti-Mouse (LiCor 926-32210, Lot# C40213-01) were then used as secondary probes agasint primary antibodies. Immuno-blots were imaged with the LiCor Odyssey^®^ CLx Imaging System.

### siRNA knockdown

Three Silencer^®^ select siRNAs against each gene target were purchased from Ambion, ThermoFisher Scientific. Information on the siRNAs may be found in Table S7. 10 nM of pool-of-three different siRNAs were used to knockdown each target gene.

### RNA immunoprecipitation Assay

2 day LIF- or RA-treated mouse ESCs were UV crosslinked at 450 mJ/cm^2^. Cells extracts were made using a lysis buffer (100 mM Tris pH 7.4, 150 mM NaCl, 1% NP40, RNase Inhibitor, Protease Inhibitory Cocktail and 1 mM DTT). 1 mg of cell extract was taken for immunoprecipitation following which extracts were pre-cleared with 50 ul Protein A/G resin (Thermo Scientific, 53132) for an hour at 4 degrees. Pre-cleared lysates were incubated with Igf2bp1 antibody (Cell Signaling Technologies, 2852S) or IgG (Cell Signaling Technologies, 3900S) control overnight at 4 degrees. Antibody was immunoprecipitated using Protein A/G coated resin and subsequently washed four times with lysis buffer. To release the RNA from the protein complex, the pull-down sample was treated with 5 mg/ml Proteinase K at 37 degrees for 10 mins. The RNA was extracted using TriZol. Small RNA libraries were made using the isolated RNA as described using TruSeq Small RNA Prep Kit. The libraries were sequenced and analysed as discussed previously. For QPCR analysis, 1ul of the isolated RNA was reverse transcribed and used in QPCR.

### Myc-Luc reporter activity assay

The HCT116 Myc-Luc reporter cells were reverse transfected in 384 well plates. Transfection mix consisting of transfectants, Lipofectamine 2000 (used according to manufacturer’s recommendations) and OptiMEM were pipetted into respective wells (20 µL per well). 5,500 cells in 30 µl of 16.7% FBS supplemented DMEM media were then seeded in each well. After 72 hrs, cell viability was assayed for using PrestoBlue reagent (ThermoFisher Scientific A13262), followed by SteadyGlo luciferase assay (Promega E2550) for Myc activity. Fluorescence and luminescence readings were obtained with the Tecan Infinite M1000 PRO Multimode Microplate Reader.

### Mouse skin isolation and FACS

Back skin of four 8 weeks CD1 mice was trypsinized overnight at 4 °C. The epidermis was scraped and chopped into fine pieces to further trypsinize to isolate single cells. The cells were washed with PBS and re-suspended in PBS+1% FBS. Cells were stained for 1 hr on ice with Integrin α6-PE (ab95703, Abcam) and/or CD34-FITC conjugated (11-0341-85, eBioscience) antibodies. Cells were sorted on FACS Aria™ (BD Biosciences) based on CD34^+^α6^+^ (Bulge stem cells), CD34^−^α6^+^ (Basal epithelia) and CD34^−^α6^−^ (differentiated keratinocytes) (Plot S1). This was followed by RNA extraction and small RNA library prep (TruSeq Small RNA Prep Kit, Illumina). Small RNA libraries were sequenced on Illumina NextSeq 500 platform.

MDA-MB-231 and HS578T breast cancer cells were sorted for CD44/CD24 expression with AlexaFlour488 conjugated α-CD44 (Stem Cell, Clone IM7, Lot #SC02931) and α-CD24 (Santa Cruz Biotechnology, FL-80, SC-11406 Lot #E1413). Cells were sorted with the FACS Aria™ Fusion Sorter (BD Biosciences) (Plot S2 and S3) operated by the A*STAR SIgN common FACS facility. Each sample was analyzed with negative controls and single channel/color controls.

## Author Contributions

DP, SR & RD conceived and designed the study. DGRY, SK performed experiments. VL, JLYK, DP performed all computational analyses. JEP & SKS contributed critically to mass spectrometry; JKC contributed to experiments on HMEC lines; JLL & MJSL helped with cell culture and Myc reporter assays; HG, JI & JMN collaborated on the Click-IT assay and RA time-point polysome profiling. IBHT & NGI contributed clinical tumor samples. DP, SR, RD, DGRY, SK wrote the manuscript with extensive input from all authors. DP, SR & RD declare no competing financial interests.

## Data Access

The sequencing data generated for this study has been deposited in Sequence Read Archives, National Center for Biotechnology Information (SRA). The raw sequences can be accessed with the project ID SRP079660.

## Acknowledgments

We would like to extend our gratitude to Drs Wan Yue (GIS), Francesc Xavier Roca Castella (NTU) and Apurva Sarin (inStem) for comments and critical review of the manuscript. We also thank Drs. Xiaoqian Zhang, and Abil Saj for help with PDx models and cell culture for RNA-seq. We are thankful for technical support from the following: The Centre for High-throughput Phenomics (CHiP-GIS) Singapore, for RNAi-based functional validation of candidate proteins identified in tsRNA pulldown screen; GIS-NGS facility and Genotypic Technology, Bangalore, for sequencing; the A*STAR SiGN Common FACS Facility, and Central Imaging and Flow Facility (CIFF, Bangalore) for FACS-based applications; and the Electron Microscopy facility at inStem, Bangalore. This work was supported by the following grants: Wellcome-DBT India Alliance Intermediate Fellowship (500160/Z/09/Z) to DP.; DBT Grant (BT/PR8655/AGR/36/759/2013) to RD, SR; inStem core funds through the DBT to SR and DP; Agency for Science, Technology and Research (A*STAR)-GIS core funds to RD. DGRY was supported by the A*STAR Graduate Academy (AGA).

